# No detectable alloreactive transcriptional responses during donor-multiplexed single-cell RNA sequencing of peripheral blood mononuclear cells

**DOI:** 10.1101/2020.02.12.946509

**Authors:** Christopher S. McGinnis, David A. Siegel, Guorui Xie, Mars Stone, Zev J. Gartner, Nadia R. Roan, Sulggi A. Lee

**Affiliations:** University of California San Francisco, Department of Pharmaceutical Chemistry, San Francisco, CA, USA; University of California San Francisco, Department of Medicine, Division of HIV/AIDS, San Francisco, CA, USA; Gladstone Institute of Virology and Immunology, San Francisco, CA, USA; Department of Urology, University of California, San Francisco, San Francisco, CA; University of California San Francisco, Department of Laboratory Medicine, San Francisco, CA, USA; Vitalant Research Institute, University of California, San Francisco, San Francisco, CA, USA; Chan Zuckerberg BioHub, University of California, San Francisco, San Francisco, CA, USA; Center for Cellular Construction, University of California, San Francisco, San Francisco, CA, USA; Helen Diller Family Comprehensive Cancer Center, San Francisco, CA, USA

## Abstract

Single-cell RNA sequencing (scRNA-seq) provides high-dimensional measurement of transcript counts in individual cells. However, high assay costs limit the study of large numbers of samples. Sample multiplexing technologies such as antibody hashing and MULTI-seq use sample-specific sequence tags to enable individual samples (e.g., different patients) to be sequenced in a pooled format before downstream computational demultiplexing. Critically, no study to date has evaluated whether the mixing of samples from different donors in this manner results in significant changes in gene expression resulting from alloreactivity (i.e., response to non-self immune antigens). The ability to demonstrate minimal to no alloreactivity is crucial to avoid confounded data analyses, particularly for cross-sectional studies evaluating changes in immunologic gene signatures,. Here, we compared the expression profiles of peripheral blood mononuclear cells (PBMCs) from a single donor with and without pooling with PBMCs isolated from other donors with different blood types. We find that there was no evidence of alloreactivity in the multiplexed samples following three distinct multiplexing workflows (antibody hashing, MULTI-seq, and *in silico* genotyping using souporcell). Moreover, we identified biases amongst antibody hashing sample classification results in this particular experimental system, as well as gene expression signatures linked to PBMC preparation method (e.g., Ficoll-Paque density gradient centrifugation with or without apheresis using Trima filtration).

## INTRODUCTION

Recent advances in single-cell RNA sequencing (scRNA-seq) technologies have dramatically increased assay throughput from ∼10^2^ to 10^4^ −10^6^ cells per experiment^1^. However, traditional applications of scRNA-seq workflows (e.g., 10x Genomics) require individual samples to be processed in parallel, which translates to prohibitively-high assay costs for population-scale studies requiring large numbers of samples. Several scRNA-seq sample multiplexing technologies have been developed which enable users to circumvent this limitation by processing samples in a pooled format^2-8^. By avoiding the usual requirement for processing distinct samples individually, these technologies increase scRNA-seq cell and sample throughput while minimizing technical confounders (e.g., doublets and batch effects). Two main types of sample multiplexing approaches have been described: (i) *in silico* genotyping using single nucleotide polymorphisms (SNPs) and (ii) tagging cell membranes with sample-specific DNA barcodes using lipid-modified oligonucleotides (LMOs; e.g., MULTI-seq)^2^ or DNA-conjugated antibodies^3,4^ (e.g., BD single-cell multiplexing kit (SCMK)^9^). Despite the increasing popularity of sample multiplexing, direct measures of transcriptional changes induced by mixing human samples from different individuals during scRNA-seq sample preparation have not been performed. Determining the extent to which these changes might occur is critical, as they would confound cross-sectional data interpretation.

Mixing-specific transcriptional responses could theoretically occur during sample preparation when peripheral blood mononuclear cells (PBMCs) from blood-type-mismatched donors are mixed together prior to scRNA-seq. It is well known that co-culturing PBMCs from donors with different blood types causes a rapid and potent allogeneic response^10-12^. During the allogeneic response, T lymphocytes are stimulated through T-cell receptor binding to ‘non-self’ major and minor histocompatibility complex proteins expressed by foreign antigen-presenting cells. These immunomodulatory interactions are triggered following *in vitro* co-culture of blood-type-mismatched immune cells and in *in vivo* contexts such as organ transplantation and graft-vs-host disease. Although samples are maintained on ice during scRNA-seq sample preparation, it is unclear whether the allogeneic response would occur at low temperatures or whether transient periods of warming (e.g., during droplet emulsion at room temperature) are sufficient to drive alloreactivity. Directly assessing whether alloreactivity will confound downstream scRNA-seq analyses is a critical benchmark for large-scale immunological studies^13^ and sample-multiplexing experiments, writ large.

Here, we performed scRNA-seq using the 10x Genomics platform on PBMC samples isolated from eight blood-type-mismatched donors pooled under conditions where cells from a single donor were processed in isolation or after donor pooling. Donor identities for each cell were assigned using SCMK and MULTI-seq data, as well as *in silico* genotyping classifications using souporcell^14^. We did not observe significant changes in global gene expression profiles linked to donor mixing. Moreover, we did not observe any statistically-significant changes in the expression of genes associated with alloreactivity in CD4+ T-cells. As a result, we conclude that pooling samples during sample preparation for 10x Genomics-based scRNA-seq does not result in any detectable alloreactivity at the RNA level.

## STUDY DESIGN

To assess whether mixing PBMC donors causes alloreactivity during scRNA-seq, we performed a cross-sectional study of PBMCs isolated from 8 healthy donors with different blood types (Fig. 1; Experimental Methods). Donors were selected based on the diversity of blood types (e.g., A, B, O, and Rhesus factor +/-) and PBMC samples were tagged with donor-specific MULTI-seq^2^ and SCMK DNA barcodes^9^. PBMCs were mixed for 30 minutes at 4°C prior to emulsion across four droplet microfluidics lanes (10x Genomics) at room temperature. We anticipated that any allogeneic response would theoretically occur during the 30-minute pooled incubation and emulsions steps. We hypothesized that if co-incubation of blood-type-mismatched PBMCs for 30 minutes at 4°C causes detectable alloreactivity, then mixed and unmixed donor A PBMCs would exhibit more variable gene expression profiles than what is observed due to technical variation.

**Figure 1:**
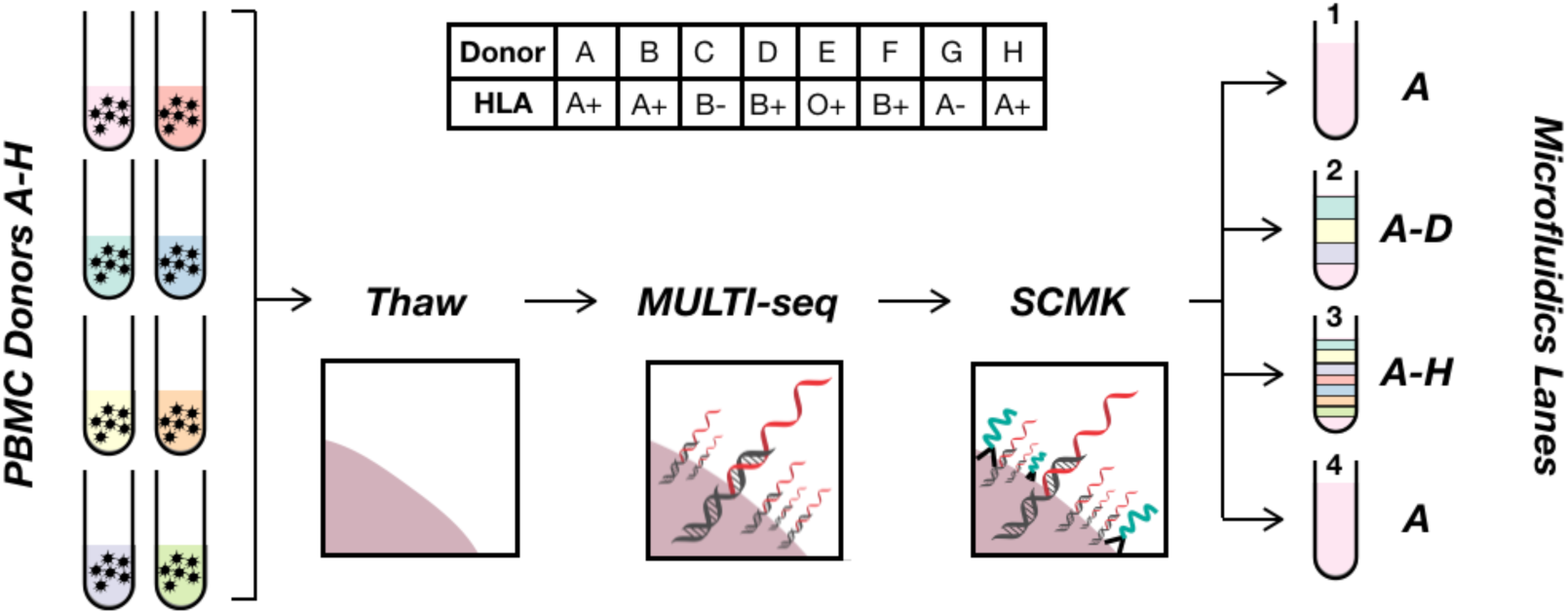
Schematic overview of experimental design. PBMCs from 8 healthy blood-type-mismatched donors (tubes on left, table on top) were barcoded with MULTI-seq LMOs (black double-helix hybridized to red DNA barcode) and BD single-cell multiplexing kit (SCMK) antibodies (black antibody conjugated to teal DNA barcode). Cells were then strategically pooled to directly assess whether mixing blood-type-mismatched PBMCs during scRNA-seq causes alloreactivity.

## RESULTS

### MULTI-seq classifies PBMCs more accurately than SCMK

To assess the performance of MULTI-seq and SCMK, we compared the results of three distinct demultiplexing workflows on donor A-H PBMCs from microfluidic lane #3: (i) deMULTIplex, (ii) demuxEM, and (iii) souporcell. deMULTIplex^2^ and demuxEM^4^ are algorithms that function on sample barcode count matrices, while souporcell is an *in silico* genotyping pipeline that functions on gene expression data^14^. MULTI-seq and SCMK classifications were largely consistent with the souporcell results (Fig. 2A) – e.g., amongst cells classified as donors A-H using souporcell, 99.9% and 99.0% of donor classifications were consistent for MULTI-seq and SCMK, respectively. However, while 1.5% of cells remained unclassified following MULTI-seq demultiplexing, 36.2% of cells remained unclassified after SCMK demultiplexing. This decrease in classification efficiency was also observed when compared to the demuxEM results (Table 1).

**Table 1:** MULTI-seq and SCMK classification performance comparison relative to *in silico* genotyping results using souporcell. MULTI-seq and SCMK classification performance was determined using two statistics: (i) % Unclassified, calculated as the proportion cells classified into donor groups by souporcell that remain unclassified by MULTI-seq or SCMK (ii) % Donor Match, calculated as the proportion of cells classified into the same donor group by MULTI-seq or SCMK and souporcell. Both metrics were computed using results from both deMULTIplex and demuxEM.

| Method | MULTI-seq | SCMK |
| --- | --- | --- |
| % Unclassified (deMULTiplex) | 1.5% | 36.2% |
| % Unclassified (demuxEM) | 2.8% | 35.4% |
| % souporecell Donor Match (deMULTiplex) | 99.9% | 99.0% |
| % souporecell Donor Match (demuxEM) | 98.8% | 97.7% |

**Figure 2:**
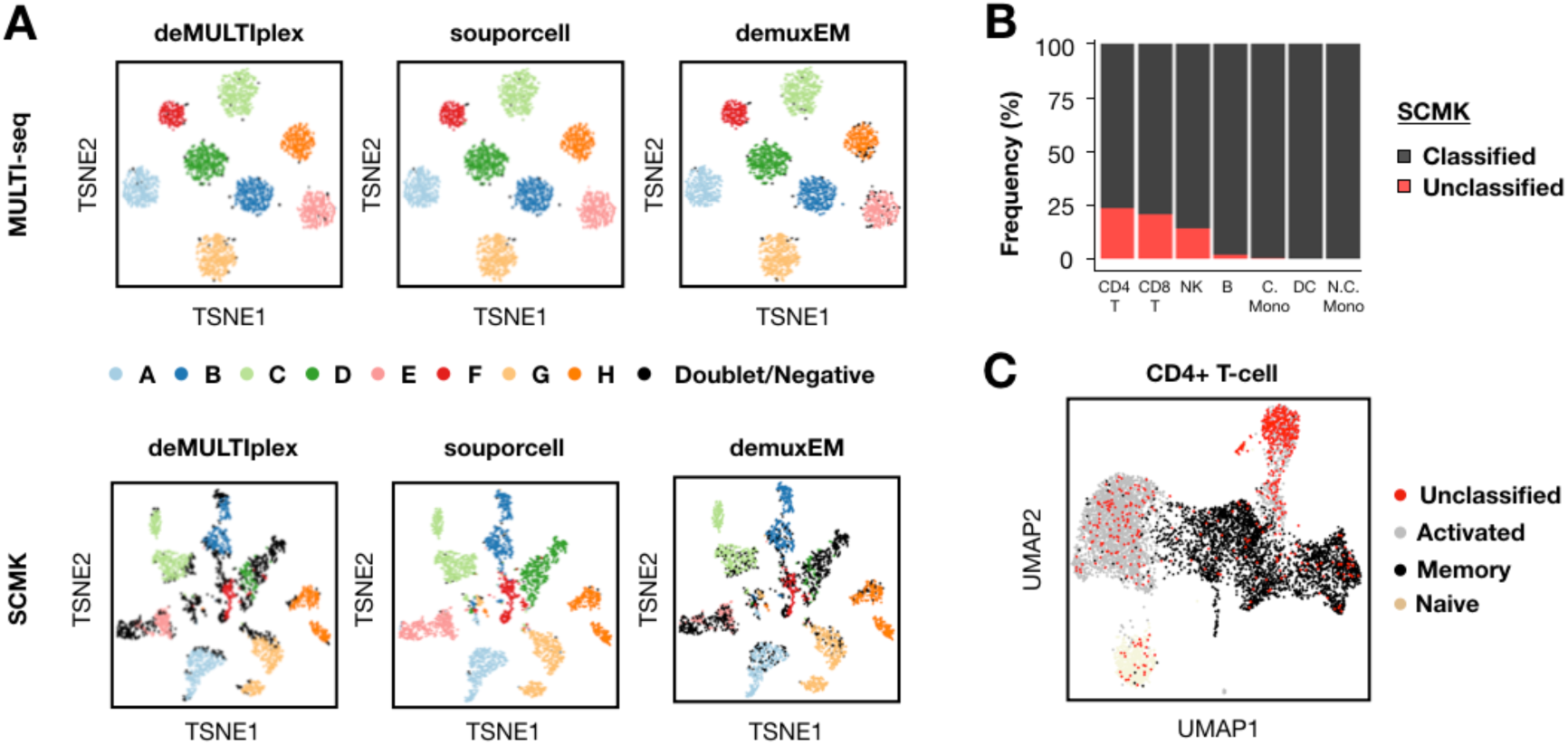
MULTI-seq and SCMK classifications largely match *in silico* genotyping, with lower SCMK classification efficiency and bias against activated CD4+ T-cells. (A) Sample classification results from three demultiplexing pipelines (e.g., deMULTIplex, souporcell, and demuxEM) projected onto MULTI-seq (top) and SCMK (bottom) sample barcode space. n = 4,032 cells from microfluidic lane #3. (B) Frequencies of classified and unclassified cells across all PBMC cell types following SCMK sample demultiplexing. (C) Localization of SCMK unclassified cells in CD4+ T-cell gene expression space. n = 6,879 CD4+ T-cells.

To assess whether cells that remained unclassified following SCMK demultiplexing were randomly distributed throughout the scRNA-seq data, we computed the frequency of unclassified cells in each PBMC cell type. This analysis revealed that T lymphocytes and NK cells were especially likely to remain unclassified in SCMK data (Fig. 2B). Moreover, activated CD4+ T-cells were particularly prominent among the unclassified CD4+ T-cells (Fig. 2C; Fig. S2A). For these reasons, MULTI-seq donor classifications were used for all subsequent gene expression analyses.

### Trima apheresis introduces biologically-relevant confounders into PBMC scRNA-seq data

The PBMCs that were used in this study came from whole blood that was processed using Ficoll-Paque density gradient centrifugation. Notably, these samples either underwent (donors D-H) or did not undergo (donors A-C) apheresis using Trima filtration, a method to enhance leukocyte yield during sample preparation^15,16^. Initial inspection of MULTI-seq donor classifications revealed that PBMCs predominantly clustered according to processing method – e.g., Trima vs. Ficoll (Fig. S3A). Upon sub-clustering classical monocytes and NK cells, we observed that Trima and Ficoll classical monocytes expressed variable levels of the histone component gene HIST1H1C, as well as two genes involved in monocyte differentiation, MNDA and CEBPB (Fig. S3B, left)^17^. Moreover, we observed that Trima and Ficoll NK cells differentially expressed the immune cytokine IFNG, cytolytic genes GZMA and PRF1, and the stress marker JUN (Fig. S3B, right)^18^. These results suggest that apheresis using Trima filters induces confounding changes in gene expression patterns associated with differentiation state, cytolytic activity, and stress across multiple PBMC cell types. These signatures are consistent with prior observations^19^ and should be accounted for in future analyses. Thus, to avoid these confounding effects when comparing donor-and mixing-specific expression profiles, we restricted our subsequent analyses to PBMC samples processed without Trima filtration.

### Mixing blood-type-mismatched PBMCs during scRNA-seq sample preparation does not cause a detectable allogeneic transcriptional response

To assess whether mixing PBMCs induces alloreactivity during multiplexed scRNA-seq sample preparation, we compared the expression profiles of mixed and unmixed donor A PBMCs. Mapping the densities of mixed and unmixed donor A sample classifications onto PBMC gene expression space (Fig. 3A, top left) did not reveal any qualitative shifts in global gene expression profiles (Fig. 3A, bottom). Notably, such shifts in classification densities were observed when including PBMCs from donors B and C (Fig. 3A, top right), suggesting that natural inter-donor variation is more pronounced than variation due to PBMC mixing.

**Figure 3:**
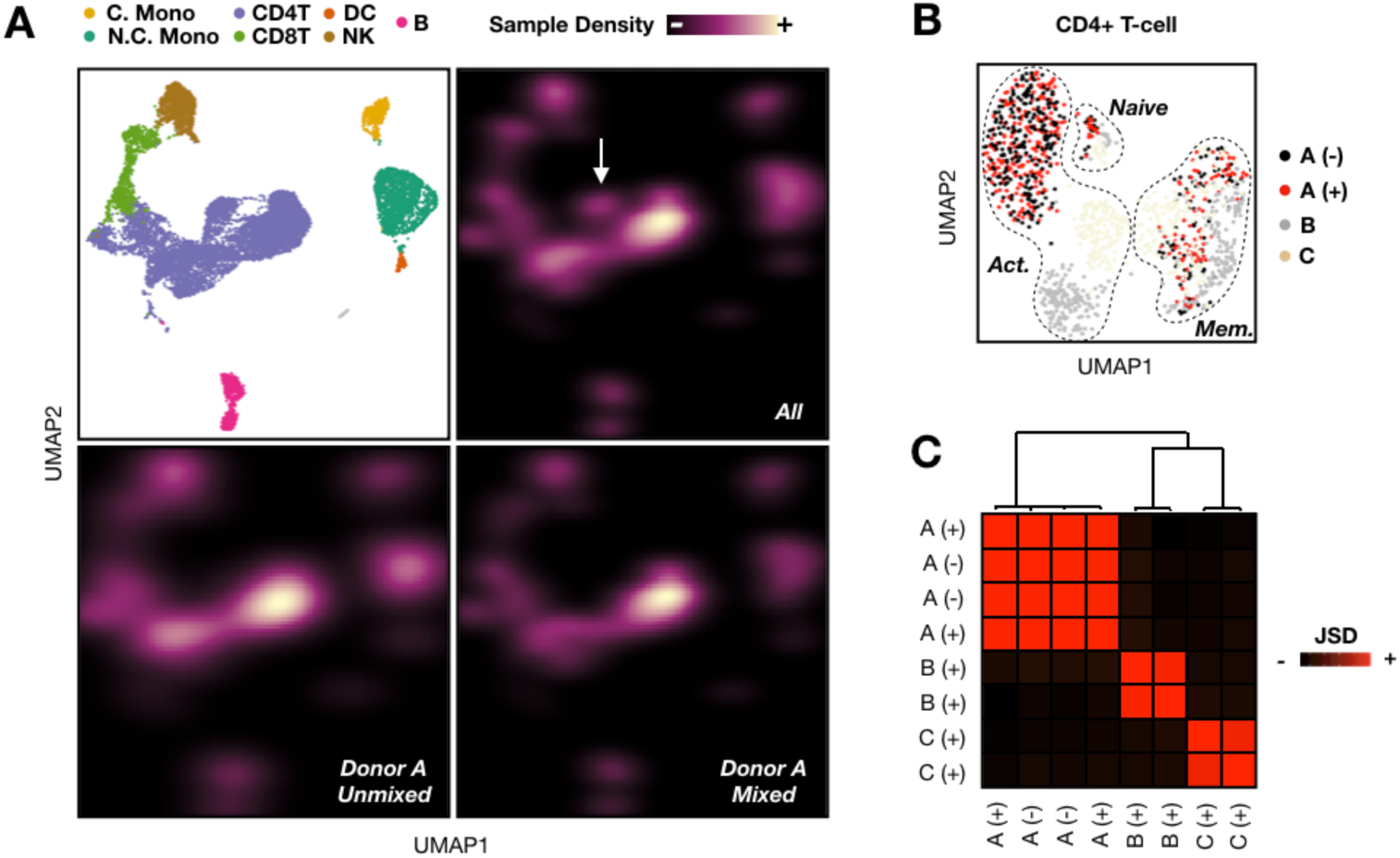
Mixing blood-type-mismatched PBMCs does not cause allogenic response during multiplexed scRNA-seq sample preparation. (A) Sample classification results plotted as densities in PBMC gene expression space (top left) grouped according to unmixed donor A PBMCs (bottom left), mixed Donor A PBMCs (bottom right), and Donors A-C PBMCs (top right). Discordant region representing donor-specific expression profiles highlighted with white arrow. (B) Evenly-subsetted CD4+ T-cell gene expression space grouped according to donor ID (e.g., A, B, C) and mixing status (e.g., - = unmixed, + = mixed). n = 1,336 CD4+ T-cells. (C) Quantitative inter-sample comparison using Jensen-Shannon Divergence (JSD) and hierarchical clustering demonstrates similarity between evenly-subsetted CD4+ T-cells from distinct healthy donors regardless of mixing.

Next, we focused on CD4+ T-cells because of their known involvement in alloreactivity^10-12^ and their relatively large prevalence in our scRNA-seq dataset. As was observed in the full dataset, mixed and unmixed donor A cells were similarly clustered together in CD4+ T-cell gene expression space (Fig. 3B). To quantify the magnitude of gene expression variability between mixed and unmixed donor A CD4+ T-cells, we used Jensen-Shannon Divergence (JSD)^20^ and hierarchical clustering to compute sample-level differences in gene expression space between the following groups: unmixed donor A, mixed donor A, donor B, and donor C (Computational Methods). This analysis reinforced the observation that CD4+ T-cells clustered predominantly by donor (average inter-donor JSD = 0.948; Fig. 3C). Moreover, we computed the average JSD due to PBMC mixing (mixed-vs-unmixed donor A JSD = 0.012) and technical noise (unmixed donor A JSD = 0.0098).

To determine whether the increased JSD for mixed CD4+ T-cells relative to technical replicates was significant, we performed a permutation test. Specifically, we reasoned that if the observed JSD differences were smaller than JSD values computed after randomly swapping donor A labels (n = 100 iterations), then gene expression variability due to mixing would be on-par with technical variability between droplet microfluidic lanes. Indeed, the average permuted JSD between donor A cells was 0.025 and was smaller than the experimental JSD only 15 times out of 100, suggesting that we could not reject the null hypothesis that differences between mixed and unmixed donor A PBMCs were on-par with technical noise (p=0.85). Similar conclusions were drawn when classical monocytes were analyzed (Fig. S2B, S2C).

Finally, we performed Gene Set Enrichment Analysis^21,22^ to determine whether certain pathways involved in immune activation and/or alloreactivity were enriched in mixed relative to unmixed donor A CD4+ T-cells. This analysis revealed that there was no statistically-significant enrichment of relevant gene sets in the mixed donor A CD4+ T-cells (Supplemental Table 1). Moreover, mixed and unmixed donor A CD4+ T-cells expressed genes that are known to be differentially expressed by lymphocytes during an allogeneic response^10-12^ at similar levels (Fig. S2D). Collectively, these results demonstrate that mixing PBMCs from different donors under multiplexed scRNA-seq sample preparation conditions does not result in a detectable allogeneic transcriptional response.

## DISCUSSION

Sample multiplexing approaches for scRNA-seq are being increasingly utilized by the single-cell genomics field to reduce assay costs while improving data breadth and quality. However, the impact of pooling distinct donor cells during scRNA-seq sample preparation on gene expression patterns has not yet been described. Here, we used the 10x Genomics scRNA-seq platform to directly compare the gene expression profiles of PBMCs prepared for sequencing alone or after mixing with blood-type-mismatched PBMCs for 30 minutes at 4°C. We found no detectable evidence of global changes in gene expression profiles (quantified using Jensen-Shannon Divergence and semi-supervised hierarchical clustering), gene signature-level differences (using Gene Set Enrichment Analysis), or following interrogation of previously-reported marker genes linked to alloreactivity. Although PBMCs actively participating in an allogeneic response were not included in this study, these results demonstrate that neither lymphocytes (e.g., CD4+ T-cells) or myeloid cells (e.g., classical monocytes) respond to mixing with blood-type-mismatched PBMCs in a fashion that is qualitatively or quantitatively distinct from technical noise.

In addition to the alloreactivity analysis, we found that donor-classified cells in the SCMK demultiplexing results were biased against activated CD4+ T-cells. This finding contrasts with the original Cell Hashing report^3^, where PBMCs were systematically demultiplexed following incubation with a panel of DNA-conjugated antibody selected for their uniform targeting of all known PBMC cell populations. Notably, the exact antigens targeted by the commercial SCMK reagents we used in this study are unknown, but our findings would suggest that the “universal” antigens targeted by the antibodies in this kit are not truly universal. Thus, users should exercise caution before using these reagents, for example by testing the uniformity of antibody binding using flow-cytometry prior to single cell library prep. In any case, validation of surface antigen expression across all cells in a given experimental system and/or careful data quality-control is necessary to avoid systematically-biased interpretations.

Collectively, this study provides support for future donor-multiplexed scRNA-seq studies of PBMCs. Our results illustrate that alloreactivity can be disregarded as a potential scRNA-seq data confounder in large-scale, cross-sectional single-cell genomics experiments. Moreover, our results demonstrate how PBMC preparation method (e.g., Ficoll-Paque density gradient centrifugation with or without Trima filtration) and sample multiplexing technology (e.g., SCMK) can introduce confounding variables into scRNA-seq data.

## EXPERIMENTAL METHODS

### scRNA-seq sample preparation

PBMCs were provided by the Vitalant Research Institute. PBMCs were thawed at 37°C and washed one time with warm media (RPMI (Corning, Cat#10-040-CV), supplemented with 10% FBS (VWR, Cat#97068-085) and Benzonase (1:1000, Sigma-Aldrich, Cat#E1014)) and one time with 2% FBS in PBS (Ca^++^ and Mg^+^ free, Corning, Cat#21-031-CV) before counting cells (Nexcelom K2). Live cells were then enriched using a dead-cell removal kit (STEM Cell, Cat#17899). Live cells were then washed with PBS and labeled with LMOs, as described previously^2^. LMOs were then quenched while washing cells with 1% BSA in cold PBS. Cells were then incubated with 5ul human Fc Block with 95ul 2% FBS in PBS at 4°C for 15 mins before staining with SCMK and AbSeq antibodies (BD Biosciences) at 4°C for 60 min. Notably, AbSeq data was not analyzed in this study. Cells were then washed twice by using 0.04% BSA (Non-acetylate, Sigma-Aldrich; B6917)) in cold media before incubation for 30 minutes at 4°C either alone (e.g., donor A) or in a pooled format (e.g., donors A-D or A-H). Finally, cells were isolated via droplet emulsion across four 10x Genomics microfluidic lanes (V2).

### Next-generation sequencing and library preparation

cDNA expression, MULTI-seq, and SCMK libraries were prepared as described previously^2^ or according to supplier recommendations. Notably, following size-selection of MULTI-seq and SCMK oligos after cDNA amplification, two separate sample-index PCRs were performed for the MULTI-seq and SCMK oligos using separate i7 indices. cDNA expression and SCMK libraries were pooled and sequenced on a single NovaSeq 6000 lane. MULTI-seq libraries were sequenced separately using the MiSeq (V3).

## COMPUTATIONAL METHODS

### scRNA-seq, MULTI-seq, and SCMK data pre-processing

Following next-generation sequencing, cDNA expression FASTQs were pre-processed and read-depth normalized using Cell Ranger (v3.0.0). FASTQs were aligned to the hg19 reference transcriptome. MULTI-seq and SCMK FASTQs were pre-processed using the ‘MULTIseq.preProcess’ function in the ‘deMULTIplex’ R package^2^. Notably, because the MULTI-seq and SCMK barcode sequences are 8 and 40 nucleotides in length, respectively, the Hamming Distance alignment threshold applied to SCMK data was increased to 5 (default = 1) to account for the increased probability of random sequencing errors.

### scRNA-seq data quality-control

Raw gene expression matrices were quality-controlled using Seurat^23,24^ (Fig. S1). First, cells with fewer than 250 RNA unique molecular identifiers (UMIs) and genes with fewer than 3 UMIs across all cells were discarded. This parsed dataset comprised of 20,353 cells and 17,908 genes was then normalized using ‘SCTransform’ prior to unsupervised clustering and dimensionality reduction using PCA and UMAP. We then removed 3,726 low-quality cells selected via membership in clusters associated with low total RNA UMIs and/or high proportions of mitochondrial gene expression (Fig. S1A).

Next, we split the cleaned dataset by lane-of-origin and applied DoubletFinder^25^ to each data subset. Notably, DoubletFinder was run on each lane independently to ensure that representative artificial doublets were constructed for each lane (e.g., multi-donor doublets were not generated for the unmixed data subsets). Moreover, we did not use MULTI-seq, SCMK, or souporcell classification results for doublet detection because each approach would produce different results for each lane (e.g., no doublets detected for single-donor datasets). DoubletFinder parameters were kept constant for each lane (e.g., pN = 0.25, pK = 0.01), resulting in the removal of 1,287 heterotypic doublets (Fig. S1B).

### PBMC cell type annotation

We annotated a final dataset of 15,340 cells using previously-established PBMC cell type marker genes^23,24^ (Fig. 1C) and identified most major cell types found in peripheral blood: CD4+ T lymphocytes (IL7R+CD8A-), CD8+ T lymphocytes (CD8A+), NK cells (SPON2+), B lymphocytes (MS4A1+), classical monocytes (CD14+), non-classical monocytes (FCGR3A+), and dendritic cells (CLEC10A+). Upon sub-clustering 6,879 CD4+ T-cells, we identified three subtypes using marker genes described previously^21,22^ (Fig. S3A): activated (SELL-lo, CREM-hi, GPR183-hi), naïve (SELL-hi, CREM-lo, GPR183-lo), and memory (SELL-hi, CREM-lo, GPR183-hi).

### Sample demultiplexing

Cells were assigned into donor groups using three different workflows. First, MULTI-seq and SCMK barcode count matrices were fed into the ‘classifyCells’ and ‘findThresh’ functions in the ‘deMULTIplex’ R package^2^. Second, MULTI-seq and SCMK barcode count matrices and the raw .h5 file (from Cell Ranger) were fed into ‘demuxEM’ (p=8), an alternative sample classification pipeline written in Python^4^. Third, position-sorted BAM files (from Cell Ranger) were fed into the *in silico* genotyping pipeline, souporcell (k=8)^13^. Notably, all methods were only applied to the 8-donor scRNA-seq data to enable robust comparisons. Upon verifying result consistency between *in silico* genotyping and MULTI-seq classification results, MULTI-seq classifications were used for all downstream analyses.

### JSD analysis

To perform global comparisons of gene expression profiles between mixed and unmixed PBMCs, we performed the following workflow. First, CD4+ T-cells and classical monocytes were computationally down-sampled to include equal numbers of cells from the following groups: unmixed donor A, mixed donor A, donor B, and donor C. Second, UMAP embeddings were computed following the pre-processing workflow described above. Third, the UMAP embedding coordinates for each PBMC group were used to compute group-wise 2-dimensional kernel density estimations with the ‘kde2d’ function in the ‘MASS’ R package^26^. Fourth, a JSD matrix representing the divergence between each sample group was computed from the kernel density estimation results using the ‘JSD’ function in the ‘philentropy’ R package^27^. Fifth, hierarchical clustering was performed on the JSD matrix using the ‘hclust’ function in the ‘stats’ R package. Differences in JSD between groups were presented after scaling from 0-1. To establish variability due to algorithm performance, the JSD calculation workflow was repeated after randomly permuting donor A classifications 100 times.

## DATA AND CODE AVAILABILITY

All code used for single cell analysis and data visualization is available via Github (github.com/chris-mcginnis-ucsf/PBMC_Allo). Submission of raw cDNA expression, MULTI-seq, and SCMK data to GEO is in process, for inquiries contact authors.

## ACKNOWLEDGEMENTS

We thank the Gladstone Genomics Core (Natasha Carli) and UCSF Center for Advanced Technology (Eric Chow) for guidance on experimental design and next-generation sequencing support. This research was supported in part by grants from the Department of Defense Breast Cancer Research Program (nos. W81XWH-10-1-1023 and W81XWH-13-1-0221), NIH (nos. U01CA199315, DP2 HD080351-01, R01AI14777, R01AI127219, R01AI143464, and 1R61DA047024), the NSF (no. MCB-1330864) and the UCSF Center for Cellular Construction (no. DBI-1548297), the 2019 Mary Anne Koda-Kimble Seed Award for Innovation, the NSF Science and Technology Center, and the amfAR Institute for HIV Cure Research (no. 109380-59-RGRL). Z.J.G. is a Chan Zuckerberg BioHub Investigator. C.S.M., is an ARCS Scholar.

## AUTHOR CONTRIBUTIONS

C.S.M., N.R.R., and S.A.L. conceptualized the study and designed experiments. C.S.M. and G.X. performed scRNA-seq experiments. C.S.M. and D.A.S. performed bioinformatics analysis. C.S.M., D.A.S., Z.J.G., N.R.R., and S.A.L. wrote the manuscript.

## SUPPLEMENTAL FIGURES

**Supplementary Figure S1:**
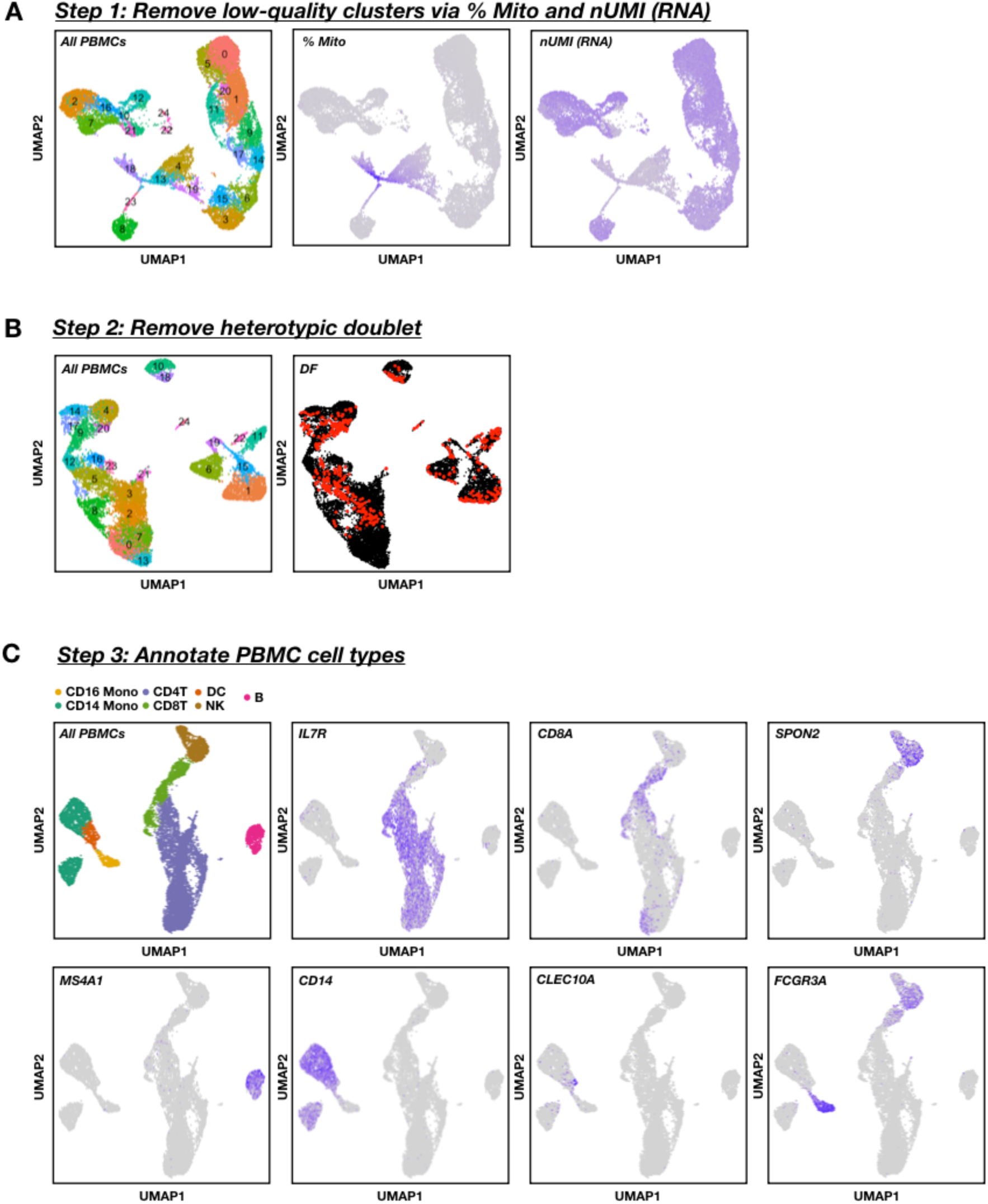
scRNA-seq data pre-processing workflow. (A) Identification of low-quality clusters in gene expression space according to percentage of mitochondrial gene expression (% Mito) and total numbers of RNA UMIs. Clusters 4, 10,13, 18-20, 23, and 24 were removed, n = 20,353 cells. (B) Identification of doublets following low-quality cell removal using DoubletFinder (DF). n = 16,627 cells. (C) Marker genes used for cell type annotations (top left) following low-quality cell and heterotypic doublet removal. n = 15,340 cells

**Supplementary Figure S2:**
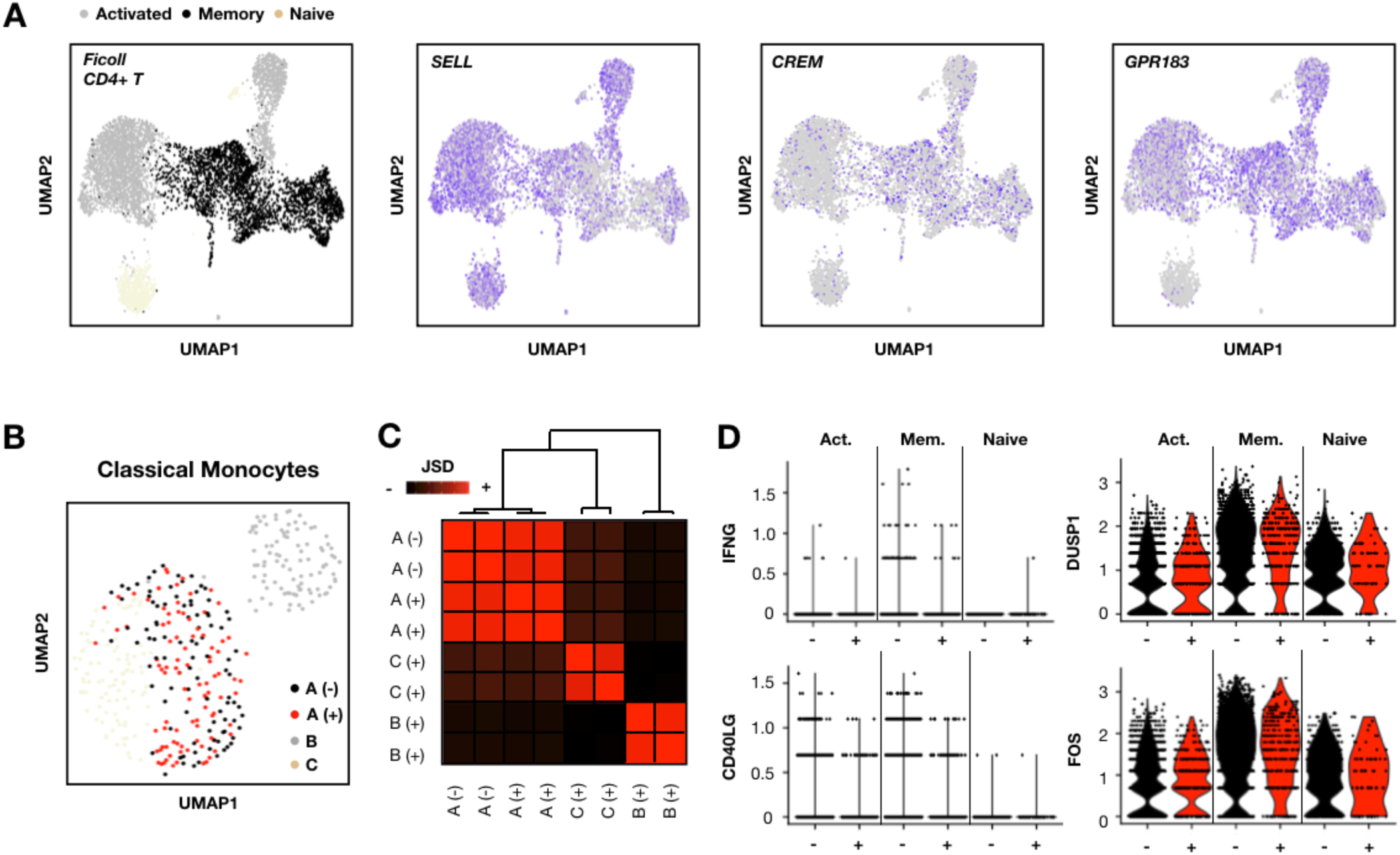
Exploring alloreactivity in CD4+ T-cell subset and classical monocytes. (A) Marker genes used for CD4+ T-cell subtype annotations (left) in Ficoll PBMCs. n = 6,879 CD4+ T-cells. (B) Evenly-subsetted classical monocyte gene expression space grouped according to donor ID (e.g., A, B, C) and mixing status (e.g., - = unmixed, + = mixed). n = 384 classical monocytes. (C) Quantitative inter-sample comparison using Jensen-Shannon Divergence (JSD) and hierarchical clustering demonstrates similarity between evenly-subsetted classical monocytes from distinct healthy donors regardless of mixing. (D) Expression of genes known to be up-regulated (e.g., IFNG and CD40LG) or down-regulated (e.g., DUSP1 and FOS) in lymphocytes during an allogenic response.

**Supplementary Figure S3:**
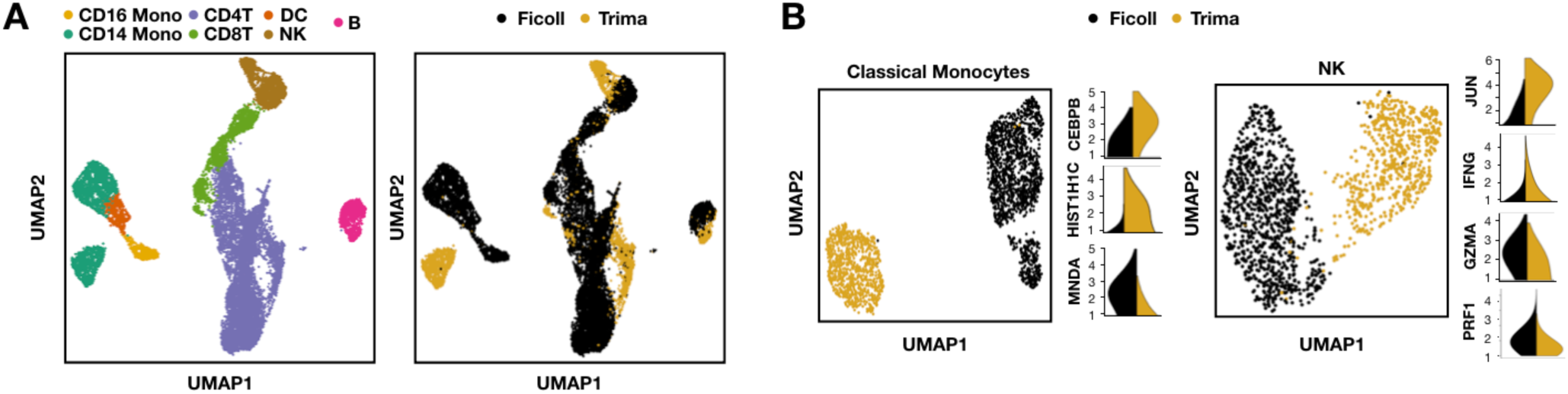
Trima-associated gene expression signatures. (A) Identification of gene expression patterns across all PBMC cell-types (left) linked to method of PBMC isolation (e.g., Ficoll or Trima; right). n = 15,340 cells. (B) Identification of Trima-specific marker genes in classical monocytes (left) and NK cells (right). n = 2,303 classical monocytes, n = 1,545 NK cells. Marker gene expression values are depicted as log1p-normalized counts.

